# Unraveling the polygenic architecture of complex traits using blood eQTL metaanalysis

**DOI:** 10.1101/447367

**Authors:** Urmo Võsa, Annique Claringbould, Harm-Jan Westra, Marc Jan Bonder, Patrick Deelen, Biao Zeng, Holger Kirsten, Ashis Saha, Roman Kreuzhuber, Silva Kasela, Natalia Pervjakova, Isabel Alvaes, Marie-Julie Fave, Mawusse Agbessi, Mark Christiansen, Rick Jansen, Ilkka Seppälä, Lin Tong, Alexander Teumer, Katharina Schramm, Gibran Hemani, Joost Verlouw, Hanieh Yaghootkar, Reyhan Sönmez, Andrew Brown, Viktorija Kukushkina, Anette Kalnapenkis, Sina Rüeger, Eleonora Porcu, Jaanika Kronberg-Guzman, Johannes Kettunen, Joseph Powell, Bernett Lee, Futao Zhang, Wibowo Arindrarto, Frank Beutner, BIOS Consortium, Harm Brugge, i2QTL Consortium, Julia Dmitreva, Mahmoud Elansary, Benjamin P. Fairfax, Michel Georges, Bastiaan T. Heijmans, Mika Kähönen, Yungil Kim, Julian C. Knight, Peter Kovacs, Knut Krohn, Shuang Li, Markus Loeffler, Urko M. Marigorta, Hailang Mei, Yukihide Momozawa, Martina Müller-Nurasyid, Matthias Nauck, Michel Nivard, Brenda Penninx, Jonathan Pritchard, Olli Raitakari, Olaf Rotzchke, Eline P. Slagboom, Coen D.A. Stehouwer, Michael Stumvoll, Patrick Sullivan, Peter A.C. ‘t Hoen, Joachim Thiery, Anke Tönjes, Jenny van Dongen, Maarten van Iterson, Jan Veldink, Uwe Völker, Cisca Wijmenga, Morris Swertz, Anand Andiappan, Grant W. Montgomery, Samuli Ripatti, Markus Perola, Zoltan Kutalik, Emmanouil Dermitzakis, Sven Bergmann, Timothy Frayling, Joyce van Meurs, Holger Prokisch, Habibul Ahsan, Brandon Pierce, Terho Lehtimäki, Dorret Boomsma, Bruce M. Psaty, Sina A. Gharib, Philip Awadalla, Lili Milani, Willem Ouwehand, Kate Downes, Oliver Stegle, Alexis Battle, Jian Yang, Peter M. Visscher, Markus Scholz, Gregory Gibson, Tõnu Esko, Lude Franke

## Abstract

**Summary:** While many disease-associated variants have been identified through genome-wide association studies, their downstream molecular consequences remain unclear.

To identify these effects, we performed *cis-* and *trans-expression* quantitative trait locus (eQTL) analysis in blood from 31,684 individuals through the eQTLGen Consortium.

We observed that *cis*-eQTLs can be detected for 88% of the studied genes, but that they have a different genetic architecture compared to disease-associated variants, limiting our ability to use *cis*-eQTLs to pinpoint causal genes within susceptibility loci.

In contrast, trans-eQTLs (detected for 37% of 10,317 studied trait-associated variants) were more informative. Multiple unlinked variants, associated to the same complex trait, often converged on trans-genes that are known to play central roles in disease etiology.

We observed the same when ascertaining the effect of polygenic scores calculated for 1,263 genome-wide association study (GWAS) traits. Expression levels of 13% of the studied genes correlated with polygenic scores, and many resulting genes are known to drive these traits.

## Main text

Expression quantitative trait loci (eQTLs) have become a common tool to interpret the regulatory mechanisms of the variants associated with complex traits through genome-wide association studies (GWAS). *cis*-eQTLs, where gene expression levels are affected by a nearby single nucleotide polymorphism (SNP) (<1 megabases; Mb), in particular, have been widely used for this purpose. However, *cis*-eQTLs from the genome tissue expression project (GTEx) explain only a modest proportion of disease heritability^1^.

In contrast, trans-eQTLs, where the SNP is located distal to the gene (>5Mb) or on other chromosomes, can provide insight into the effects of a single variant on many genes. *Trans*-eQTLs identified before^1–7^ have already been used to identify putative key driver genes that contribute to disease^8^. However, *trans*-eQTL effects are generally much weaker than those of *cis*-eQTLs, requiring a larger sample size for detection.

While *trans*-eQTLs are useful for the identification of the downstream effects of a single variant, a different approach is required to determine the combined consequences of trait-associated variants. Polygenic scores (PGS) have been recently applied to sum genome-wide risk for several diseases and likely will improve clinical care^9,10^. However, the exact consequences of different PGS at the molecular level, and thus the contexts in which a polygenic effects manifest themselves, are largely unknown. Here, we systematically investigate *trans*-eQTLs as well as associations between PGS and gene expression (expression quantitative trait score, eQTS) to determine how genetic effects influence and converge on genes and pathways that are important for complex traits.

To maximize the statistical power to detect eQTL and eQTS effects, we performed a large-scale meta-analysis in 31,684 blood samples from 37 cohorts (assayed using three gene expression platforms) in the context of the eQTLGen Consortium. This allowed us to identify significant *cis*-eQTLs for 16,989 genes, *trans*-eQTLs for 6,298 genes and eQTS effects for 2,568 genes (Figure 1A), revealing complex regulatory effects of trait-associated variants. We combine these results with additional data layers and highlight a number of examples where we leverage this resource to infer novel biological insights into mechanisms of complex traits. We hypothesize that analyses identifying genes further downstream are more cell-type specific and more relevant for understanding disease (**Figure 1B**).

**Figure 1.**
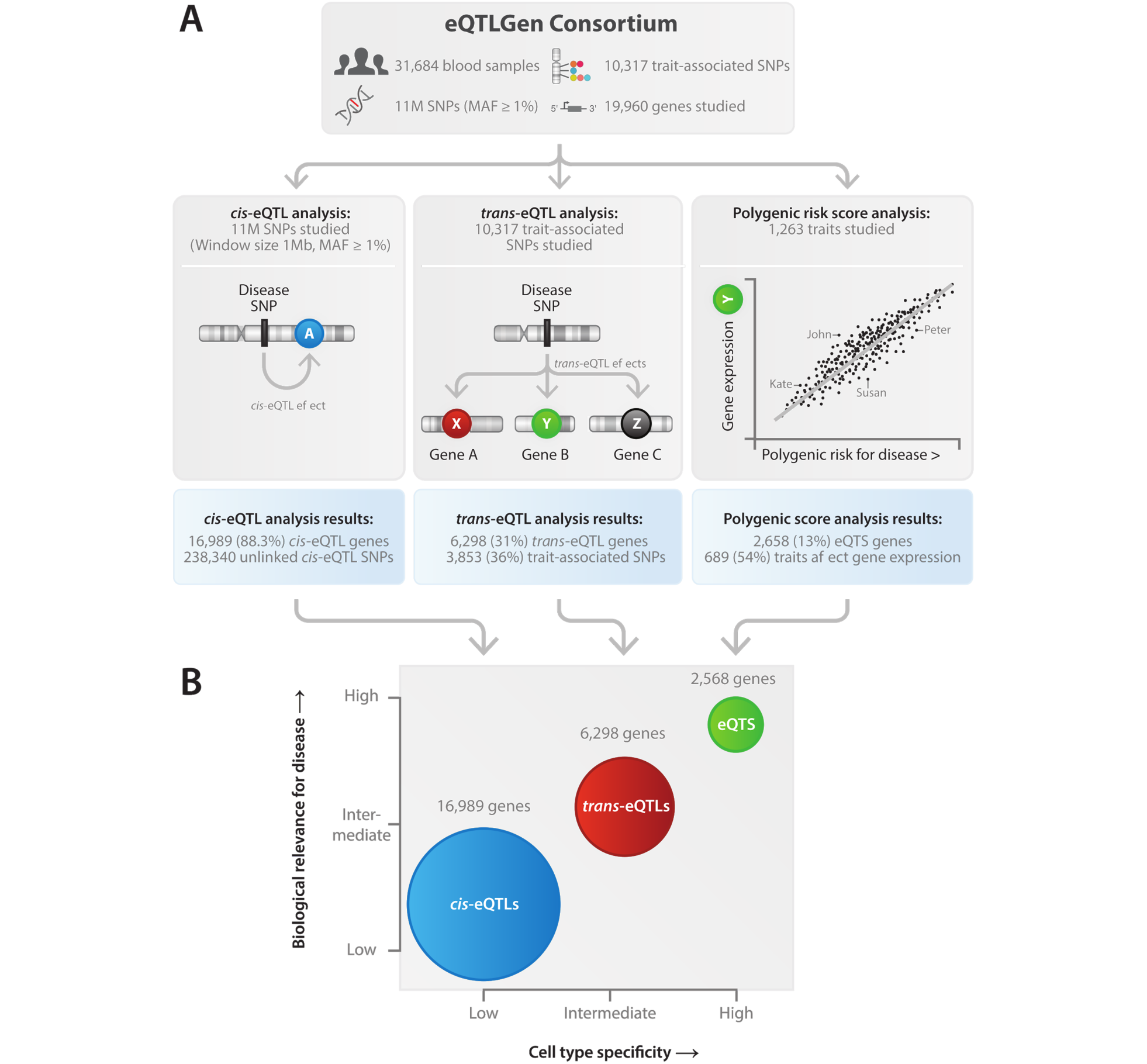
Overview of the study. (**A**) Overview of main analyses and their results. (**B**) Model of genetic effects on gene expression. *cis*-eQTL are common and widely replicable in different tissues and cell types, whereas *trans*-eQTLs and eQTS are more cell type specific. The biological insight derived from our *cis-*eQTL results are usually not well interpretable in the context of complex traits, suggesting that weaker distal effects give additional insight about biological mechanisms leading to complex traits.

## Local genetic effects on gene expression in blood are widespread and replicable in other tissues

Using eQTLGen consortium data from 31,684 individuals, we performed *cis*-eQTL, *trans*-eQTL and eQTS meta-analyses (**Figure 1A, Supplementary Table 1**). Different expression profiling platforms were integrated using a data-driven method (**Online Methods**). To ensure the robustness of the identified eQTLs, we performed eQTL discovery per platform and replicated resulting eQTLs in the other platforms, observing excellent replication rates and consistency of allelic directions (**Online Methods, Supplementary Note, Extended Data Figure 1A-C**). We identified significant *cis*-eQTLs (SNP-gene distance <1Mb, gene-level False Discovery Rate (FDR)<0.05; **Online Methods**) for 16,989 unique genes (88.3% of autosomal genes expressed in blood and tested in *cis*-eQTL analysis; **Figure 1A**). Out of 10,317 trait-associated SNPs tested, 1,568 (15.2%) were in high linkage disequilibrium (LD) with the lead eQTL SNP showing the strongest association for a *cis*-eQTL gene, (R^2^>0.8; 1kG p1v3 EUR; **Supplementary Table 2; Online Methods**). Genes highly expressed in blood but not under any detectable *cis*-eQTL effect were more likely (P=2<10^-6^; Wilcoxon two-sided test; **Figure 2A**) to be intolerant to loss-of-function mutations in their coding region^11^, suggesting that eQTLs on such gene would interfere with the normal functioning of the organism.

**Figure 2.**
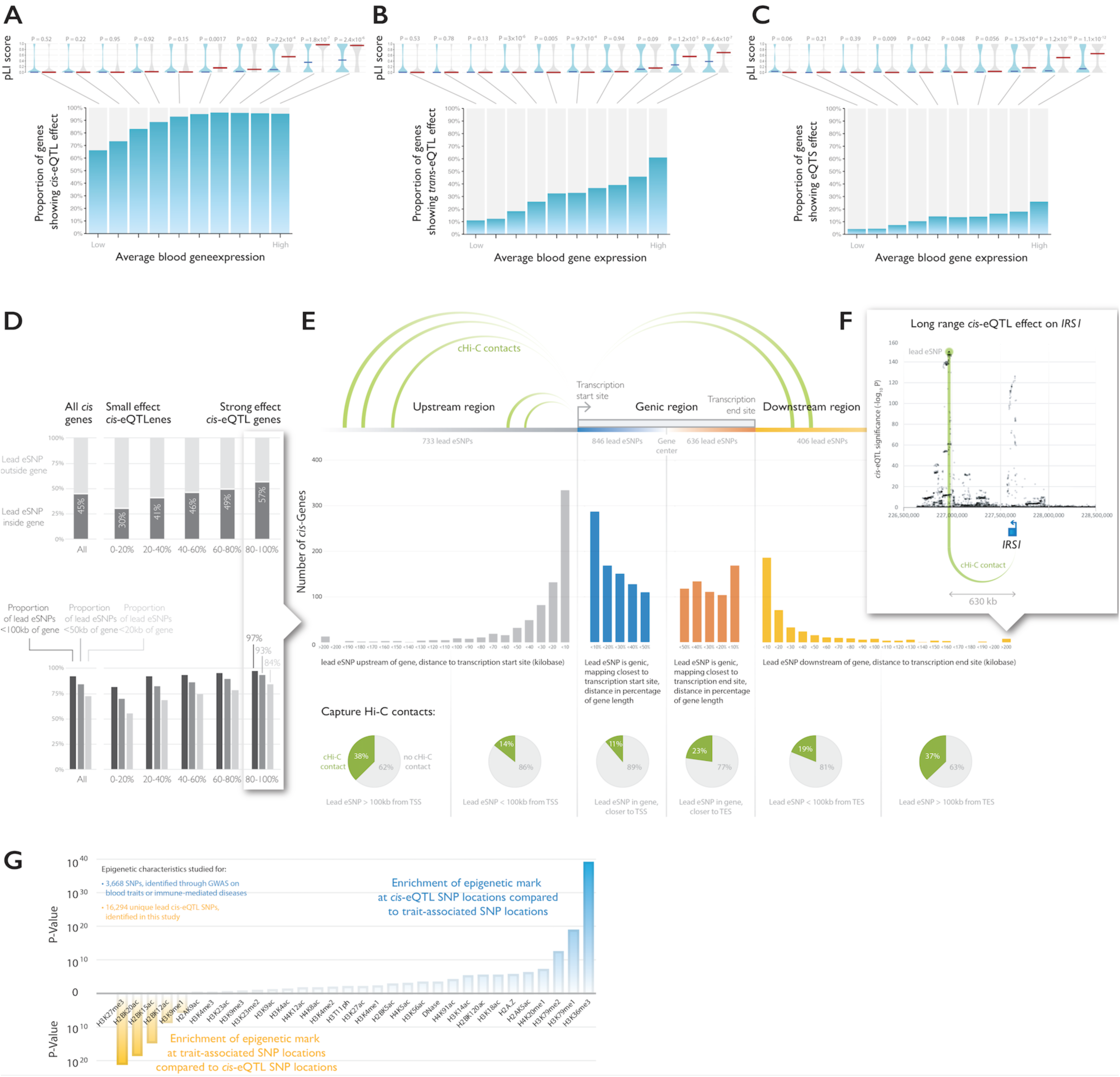
Results of the*cis-* and *trans*-eQTL analysis. All genes tested in (**A**) cvs-eQTL analysis, (**B**) *trans*-eQTL analysis, and (**C**) eQTS analysis were divided into 10 bins based on average expression levels of the genes in blood. Highly expressed genes without any eQTL effect (grey bars) were less tolerant to loss-of-function variants (Wilcoxon test on pLI scores). Indicated are medians per bin. (**D**) Genes with strong effect sizes are more likely to have a lead SNP within (top panel) or close to the gene (bottom panel) (**E**) Top *cis*-eQTL SNPs positioning further from transcription start site (TSS) and transcription end site (TES) are more likely to overlap capture Hi-C contacts with TSS. (**F**) Enrichment analyses on epigenetic marks of *cis*-eQTL lead SNPs, compared to SNPs identified through GWAS and associated to blood-related or immune-mediated diseases, reveal significant differences in epigenetic characteristics.

We observed that 92% of the lead *cis*-eQTL SNPs map within 100kb of the gene (**Figure 2D**), and this increased to 97.2% when only looking at the 20% of the genes with the strongest lead *cis*-eQTL effects. Of these strong *cis*-eQTLs, 84.1% of the lead eQTL SNPs map within 20kb of the gene. GWAS simulations^12^ indicate that lead GWAS signals map within 33.5kb from the causal variant in 80% of cases, which suggests that our top SNPs usually tag causal variants that map directly into either the promoter region, the transcription start site (TSS), the gene body, or the transcription end site (TES). For strong *cis*-eQTLs we observed that lead *cis*-eQTL SNPs located >100kb from the TSS or TES overlap capture Hi-C contacts (37%; **Figure 2E**) more often than short-range cis-eQTL effects (16%; Chi^2^ test P = 2×10^-5^), indicating that, for long-range *cis*-eQTLs the SNP and gene often physically interact to cause the *cis*-eQTL effect. For instance, a capture Hi-C contact for *IRS1* overlapped the lead eQTL SNP, mapping 630kb downstream from *IRS1* (**Figure 2F**).

We observed that our sample-size improved fine-mapping: for 5,440 protein-coding *cis*-eQTL genes that we had previously identified in 5,311 samples^1^ we now observe that the lead SNP typically map closer to the *cis*-eQTL gene (**Extended Data Figure 4**).

*Cis*-eQTLs showed directional consistency across tissues: in 47 postmortem tissues (GTEx v7^13^) we observed an average of 14.8% replication rate (replication FDR<;0.05 in GTEx; median 15.1%; range 3.6-29.7%; whole blood tissue excluded) and on average a 95.0% concordance in allelic directions (median 95.3%, range 86.7-99.3%; whole blood tissue excluded) among the *cis*-eQTLs that significantly replicated in GTEx (**Extended Data Figure 5, Supplementary Note and Supplementary Table 3**).

However, our lead *cis*-eQTL SNPs show significantly different epigenetic histone mark characteristics, as compared to 3,668 SNPs identified in GWAS (and associated to blood related traits or immune-mediated diseases to minimize potential confounding). We observed significant differences for 20 out of 32 tested histone marks with H3K36me3, H3K27me3, H3K79me1 and H2BK20ac showing the strongest difference (Wilcoxon P = 10^-39^, 10^-21^, 10^-19^ and 10^-18^,respectively), suggesting that *cis*-eQTLs have a different genetic architecture, as compared to complex traits and diseases.

We tested this for 16 well-powered complex traits (Supplementary Table 20) and observed that genes prioritized by combining *cis*-eQTL and GWAS data using summary statistics based Mendelian randomization (SMR^14^; **Online Methods**) did not overlap significantly more with genes prioritized through an alternative method (DEPICT) that does not use any *cis*-eQTL information^15^. While the genes prioritized with SMR were informative, and enriched for relevant pathways for several immune traits (**Supplementary Table 20**), non-blood-trait-prioritized genes were difficult to interpret in the context of disease. Moreover, the lack of enriched overlap between DEPICT and SMR indicates that employing *cis*-eQTL information does not necessarily clarify which genes are causal for a given susceptibility locus. As such, some caution is warranted when using a single *cis*-eQTL repository for interpretation of GWAS.

## One third of trait-associated variants have *trans-eQTL* effects

An alternative strategy for gaining insight into the molecular functional consequences of disease-associated genetic variants is to ascertain *trans*-eQTL effects. We tested 10,317 trait-associated SNPs (P ≤ 5×10^-8^; **Online Methods, Supplementary Table 2**) for *trans*-eQTL effects (SNP-gene distance >5Mb, FDR ≤ 0.05) to better understand their downstream consequences. We identified a total of 59,786 significant *trans*-eQTLs (FDR<;0.05; **Supplementary Table 4, Extended Data Figure 6**), representing 3,853 unique SNPs (37% of tested GWAS SNPs) and 6,298 unique genes (32% of tested genes; **Figure 1A**). When compared to the previous largest *trans*-eQTL metaanalysis^1^ (N=5,311; 8% of trait-associated SNPs with a significant *trans*-eQTL), these results indicate that a large sample size is critical for identifying downstream effects. Colocalization analyses in a subset of samples (n=4,339; **Supplementary Note**) using COLOC^16^ estimated that 52% of *trans*-eQTL signals colocalize with at least one *cis*-eQTL signal (posterior probability > 0.8; **Extended Data Figure 7A-B**). Corresponding colocalizing *cis*-eQTL genes were enriched for transcription factor activity (“regulation of transcription from RNA polymerase II promoter”; P <; 1.3<10^-9^**; Extended Data Figure 7C**). Finally, highly expressed genes without a detectable *trans*-eQTL effect were more likely to be intolerant to loss-of-function variants (P=6.4<10^-7^; Wilcoxon test, **Figure 2B**), similar to what we observed for *cis*-eQTLs.

In order to study the biological nature of the *trans*-eQTLs we identified, we conducted several enrichment analyses (**Supplementary Note, Extended Data Figure 8, Figure 3**). We observed 2.2 fold enrichment for known transcription factor (TF) - target gene pairs^17^ (Fisher’s exact test P = 10^-62^; **Supplementary Note**), with the fold enrichment increasing to 3.2 (Fisher’s exact test P <; 10^-300^) when co-expressed genes were included to TF targets. Those genes are potentially further downstream of respective TF targets in the molecular network. Similarly, we observed 1.19 fold enrichment of protein-protein interactions^18^ among *trans*-eQTL gene-gene pairs (Fisher’s exact test P=0.05). Some of these *cis-trans* gene pairs encode subunits of the same protein complex (e.g. *POLR3H* and *POLR1C*). While significant *cis-trans* gene pairs were enriched for gene pairs showing co-expression (Pearson R > 0.4; Fisher’s exact test P=10^-35^), we did not observe any enrichment of chromatin-chromatin contacts^19^ (0.99 fold enrichment; Fisher’s exact test P=0.3). Using the subset of 3,831 samples from BIOS, we also ascertained whether the *trans*-eQTL effect was mediated through a gene that mapped within 100kb from the *trans*-eSNP (i.e. using the *cis-*gene as G × E term). We observed significant interaction effects for 523 SNP-*cis*-trans-gene combinations (FDR <; 0.05; **Supplementary Table 5**), reflecting a 5.3 fold enrichment compared to what is expected by chance (Fisher’s exact P = 7 χ 10^-67^). For instance, for rs7045087 (associated to red blood cell counts) we observed that the expression of interferon gene *DDX58* (mapping 38bp downstream from rs7045087) significantly interacted with *trans*-eQTL effects on interferon genes *HERC5, OAS1, OAS3, MX1, IFIT1, IFIT2, IFIT5, IFI44, IFI44L, RSAD2* and *SAMD9* (**Extended Data Figure 9**).

**Figure 3.**
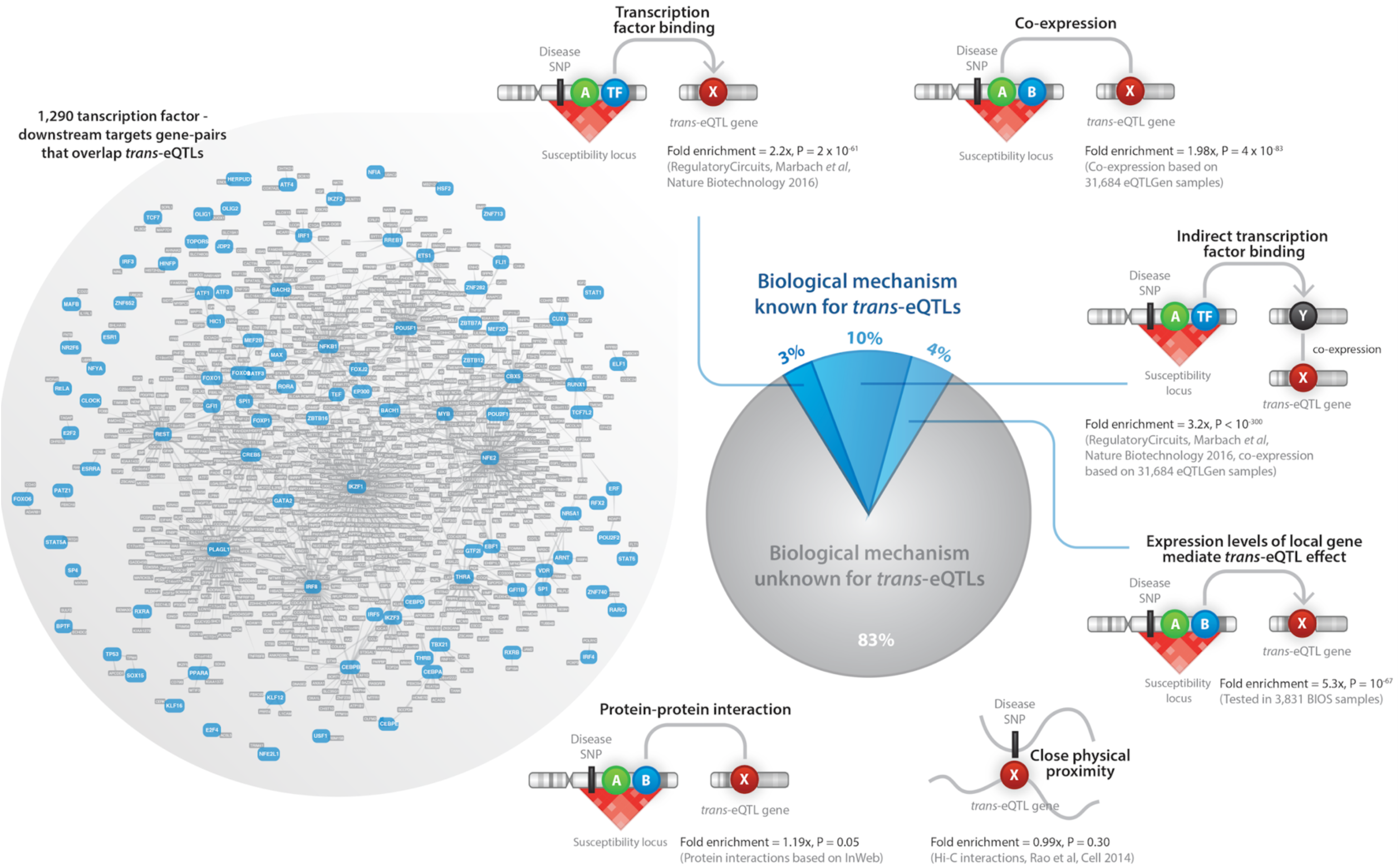
Mechanisms leading to *trans-eQTLs.* Shown are the results of enrichment analyses for known TF associations, HiC contacts, protein-protein interactions, gene co-expression and mediation analyses.

We estimate that 17.4% of the identified *trans*-eQTLs are explainable by (indirect) TF binding or mediation by *cis*-genes (**Supplementary Note**). This leaves 82.6% of the observed *trans*-eQTL effects unexplained. While it is likely that many of these *trans*-eQTLs reflect unknown (indirect) effects of TFs, we speculate that novel and unknown regulatory mechanisms could also play a role. By making all *trans*-eQTL results (irrespective of their statistical significance) publicly available, we envision this dataset will help to yield such insight in the future.

To estimate the proportion of loci where the trait-associated SNP explained the *trans*-eQTL signal in the locus, we performed locus-wide conditional *trans*-eQTL analysis in a subset of 4,339 samples for 12,991 *trans*-eQTL loci (**Online Methods; Extended Data Figure 10; Supplementary Table 6**). In 43% of these loci, we observed that the trait-associated SNP was in high LD with the *trans*-eQTL SNP having the strongest association in the locus (R^2^ > 0.8, 1kG p1v3 EUR**; Supplementary Table 7**). For 95 cases, the strongest *cis-* and *trans*-eQTL SNPs were both in high LD with GWAS SNP (R^2^ > 0.8 between top SNPs, 1kG p1v3 EUR**; Supplementary Table 7**).

The majority (64%) of *trans*-eQTL SNPs have previously been associated with blood composition phenotypes, such as platelet count, white blood cell count and mean corpuscular volume^20^. In comparison, blood cell composition SNPs from the same study comprised only 20.7% od all the tested trait-associated SNPs. This was expected, since SNPs that regulate the abundance of a specific blood cell type would result in *trans*-eQTL effects on genes, specifically expressed in that cell type.

Therefore, we aimed to distinguish *trans-eQTLs* caused by intracellular molecular mechanisms from blood cell type QTLs using eQTL data from lymphoblastoid cell line (LCL), induced pluripotent cells (iPSCs), several purified blood cell types (CD4+, CD8+, CD14+, CD15+, CD19+, monocytes and platelets) and blood DNA methylation QTL data. In total, 3,853 (6.4%) of *trans*-eQTLs showed significant replication in at least one cell type or in the methylation data (**Extended Data Figure 11, Supplementary Table 11A**). While this set of *trans*-eQTLs (denoted as the “intracellular eQTLs”) is less likely to be driven by cell type composition, we acknowledge that the limited sample size of the available *trans*-eQTL replication datasets make our replication effort very conservative. Furthermore, *trans*-eQTLs caused by variants associated with cell type proportions may be informative for understanding the biology of a trait. Therefore, we did not remove these kinds of *trans*-eQTLs from our interpretative analyses.

Next, we aimed to replicate the identified *trans*-eQTLs in the tissues from GTEx^13^. Although the replication rate was very low (0-0.03% of *trans*-eQTLs replicated in non-blood tissues, FDR <0.05, same allelic direction; **Supplementary Table 11B**), we did observe an inflation of signal (median chi-squared statistic) for identified *trans*-eQTLs in several GTEx tissues (**Extended Data Figure 12**). Non-blood tissues showing the strongest inflation were liver, heart atrial appendage and nonsun-exposed skin.

## *Trans-eQTLs* are effective for discerning the genetic basis of complex traits

As described above, *trans*-eQTLs can arise due to *cis*-eQTL effects on TFs, whose target genes show *trans*-eQTL effects. We describe below such examples, but also highlight *trans*-eQTLs where the eQTL SNP works through a different mechanism.

### Combining *cis-* and *trans*-eQTL effects can pinpoint the genes acting as drivers of *trans*-eQTL effects

For example, the age-of-menarche-associated SNP rs1532331^21^ is in high LD with the top *cis*-eQTL effect for transcription factor *ZNF131* (R^2^ > 0.8, 1kG p1v3 EUR). *cis*-eQTL and *trans*-eQTL effects for this locus co-localized for 25 out of the 75 downstream genes (**Figure 4A**). In a recent short hairpin RNA knockdown experiment of *ZNF131*^22^, three separate cell isolates showed downregulation of four genes that we identified as *trans*-eQTL genes: *HAUS5, TMEM237, MIF4GD* and *AASDH* (**Figure 4A**). *ZNF131* has been hypothesized to inhibit estrogen signaling^23^, which may explain how the SNP in this locus contributes to altering the age of menarche.

**Figure 4.**
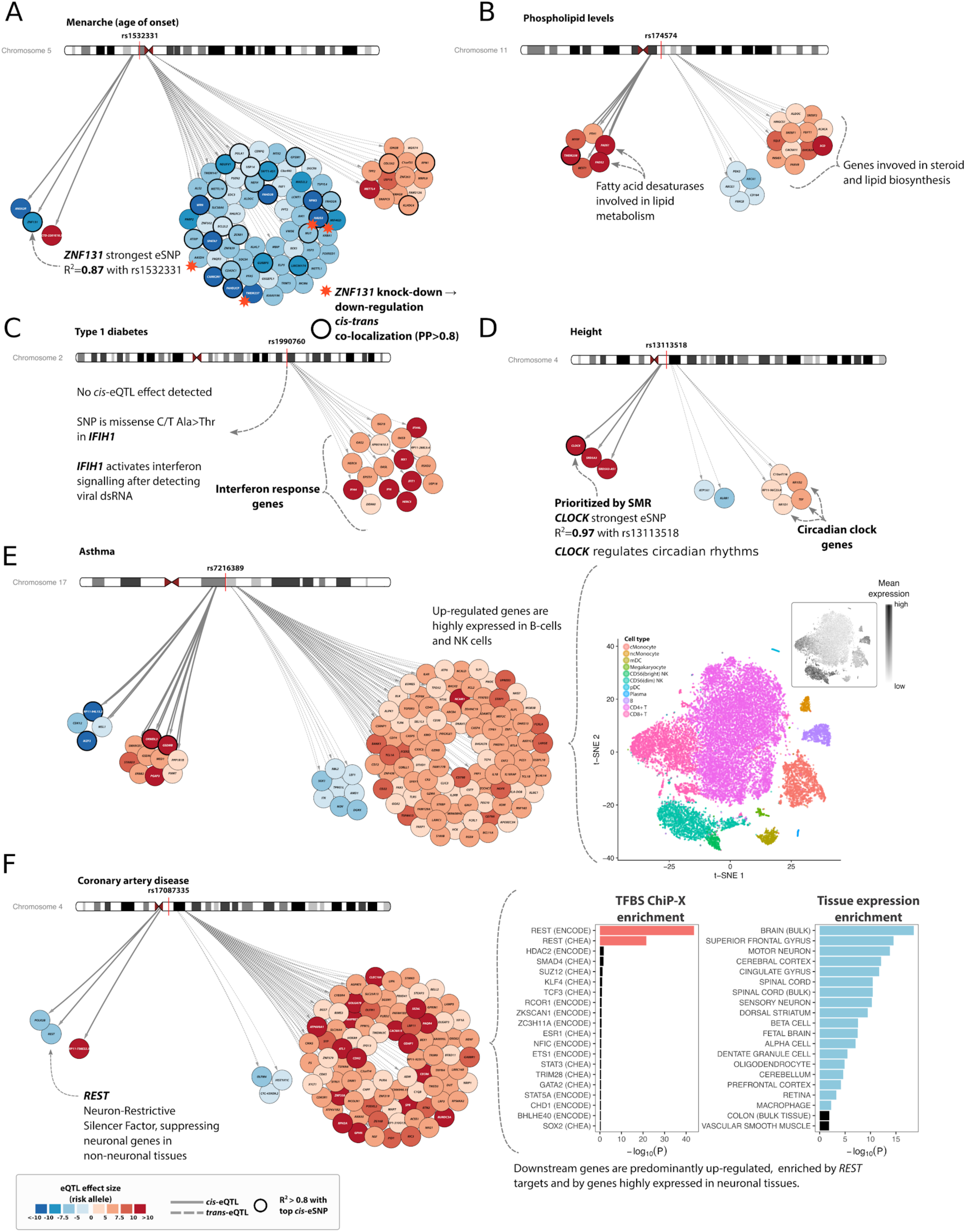
Examples of *cis-* and *trans*-eQTLs. (**A**) *cis*-eQTL on *ZNF131* is prioritized because several *trans*-eQTL genes are down-regulated by *ZNF131* in functional study**. (B**) Phospholipid-associated SNP shows *cis-and trans-eQTLs* on lipid metabolism genes. (**C**) Type I diabetes associated SNP has no *cis*-eQTLs, but *trans*-eQTL genes point to interferon signaling pathway. (**D**) Circadian rhythm genes *CLOCK* (in *cis)* and *NR1D1, NR1D2, TEF* (in *trans)* identified for height associated SNP. (**E**) eQTLs for asthma SNP tag cell type abundance of B and NK cells. (**F**) *Trans*-eQTL genes for *REST* locus are highly enriched for REST transcription factor targets and for neuronal expression.

### *Trans*-eQTLs extend insight for loci with multiple *cis*-eQTL effects

In the *FADS1/FADS2* locus, rs174574 is associated with lipid levels^24^ and affects 17 genes in *trans* (**Figure 4B**). The strongest *cis*-eQTLs modulate the expression of *FADS1, FADS2* and *TMEM258*, with latter being in high LD with GWAS SNP (R^2^>0.8, 1kG p1v3 EUR). *FADS1* and *FADS2* have been implicated^24^ since they regulate fatty acid synthesis, and consistent with their function, *trans*-eQTL genes from this locus are highly enriched for triglyceride metabolism (P < 4.1×10^-9^, GeneNetwork^25^ REACTOME pathway enrichment). Since this locus has extensive LD, variant and gene prioritization is difficult: conditional analyses in 4,339 sample subset showed that each of *cis*-eQTL gene is influenced by more than one SNP, but none of these are in high LD with rs174574 (R^2^ < 0.8, 1kG p1v3, EUR). As such, our *trans*-eQTL analysis results are informative for implicating *FADS1* and *FADS2*, whereas *cis*-eQTLs are not.

### *Trans*-eQTLs can shed light on loci with no detectable *cis*-eQTLs

Rs1990760 is associated with multiple immune-related traits (Type 1 Diabetes (T1D), Inflammatory bowel disease (IBD), Systemic Lupus Erythematosis (SLE) and psoriasis^26-29^). For this SNP we identified 17 *trans*-eQTL effects, but no detectable gene-level *cis*-eQTLs in blood (**Figure 4C**) and GTEx. However, the risk allele for this SNP causes an Ala946Thr amino acid change in the RIG-1 regulatory domain of MDA5 (encoded by *IFIH1 -* Interferon Induced With Helicase C Domain 1), outlining one possible mechanism leading to the observed *trans*-eQTLs. MDA5 acts as a sensor for viral double-stranded RNA, activating interferon I signalling among other antiviral responses. All the *trans*-eQTL genes were up-regulated relative to risk allel
e to T1D, and 9 (52%) are known to be involved in interferon signaling (**Supplementary Table 12**).

### *Trans*-eQTLs can reveal cell type composition effects of the trait-associated SNP

*Trans*-eQTL effects can also show up as a consequence of a SNP that alters cell-type composition. For example, the asthma-associated SNP rs7216389^30^ has 14 *cis*-eQTL effects, most notably on *IKZF3, GSDMB*, and *ORMDL3* (**Figure 4E**). SMR prioritized all three *cis*-genes equally (**Extended Data Figure 13**), making it difficult to draw biological conclusions (similar as we observed for the *FADS* locus). However, 94 out of the 104 *trans*-eQTL genes were up-regulated by the risk allele for rs7216389 and were mostly expressed in B cells and natural killer cells^31^ (**Figure 4E**). *IKZF3* is part of the Ikaros transcription factor family that regulates B-cell proliferation^31,32^, suggesting that a decrease of *IKZF3* leads to an increased number of B cells and concurrent *trans*-eQTL effects caused by cell-type composition differences.

### Some *trans*-eQTLs influence genes strongly expressed in tissues other than blood

We observed *trans*-eQTL effects on genes that are hardly expressed in blood, indicating that our *trans*-eQTL effects are informative for non-blood related traits as well: rs17087335, which is associated with coronary artery disease^33^, affects the expression of 88 genes in *trans* (**Figure 4F**), that are highly expressed in brain (hypergeometric test, ARCHS4 database, q-value = 2.58<10^-17^; **Figure 4F, Supplementary Table 13**), but show very low expression in blood. SNPs linked with rs17087335 (R^2^ > 0.8, 1kG p1v3 EUR) are associated with height (rs2227901, rs3733309 and rs17081935)^34,35^, and platelet count (rs7665147)^20^. The minor alleles of these SNPs downregulate the nearby gene *REST* (RE-1 silencing transcription factor), although none of these variants is in LD (R^2^<0.2, 1kG p1v3 EUR) with the lead *cis*-eQTL SNP for *REST. REST* is a TF that downregulates the expression of neuronal genes in non-neuronal tissues^36,37^. It also regulates the differentiation of vascular smooth muscles, and is thereby associated with coronary phenotypes^38^. 85 out of 88 (96.6%) of the *trans*-eQTL genes were upregulated relative to the minor allele and were strongly enriched by transcription factor targets of REST (hypergeometric test for ENCODE REST ChIP-seq, q-value = 1.36×10^-42^, **Figure 4F**). As such, *trans*-eQTL effects on neuronal genes implicate *REST* as the causal gene in this locus.

### *Trans*-eQTLs identify pathways not previously associated with a phenotype

Some *trans*-eQTLs suggest the involvement of pathways which are not previously thought to play a role for certain complex traits: SMR analysis prioritized *CLOCK* as a potential causal gene in the height-associated locus on chr 4q12 (P_**SMR**_=3×10^-25^; P_**HEIDI**_=0.02; **Figure 4D**). In line with that, height-associated SNP rs13113518^34^ is also in high LD (R^2^>0.8, 1kG p1v3 EUR) with the top *cis*-eQTL SNP for *CLOCK.* The upregulated TF CLOCK forms a heterodimer with TF BMAL1, and the resulting protein complex regulates circadian rhythm^39^. Three known circadian rhythm *trans*-eQTL genes *(TEF, NR1D1* and *NR1D2)* showed increased expression for the trait-increasing allele, suggesting a possible mechanism for the observed *trans*-eQTLs through binding of CLOCK:BMAL1. *TEF* is a D-box binding TF whose gene expression in liver and kidney is dependent on the core circadian oscillator and it regulates amino acid metabolism, fatty acid metabolism and xenobiotic detoxification (Gachon et al., 2006). *NR1D1* and *NR1D2* encode the transcriptional repressors Rev-ErbA alpha and beta, respectively, and form a negative feedback loop to suppress *BMAL1* expression^40^. *NR1D1* and *NR1D2* have been reported to be associated with osteoblast and osteoclast functions^41^, revealing a possible link between circadian clock genes and height.

### Unlinked trait-associated SNPs converge on the same downstream genes in *trans*

We subsequently ascertained, per trait, whether unlinked trait-associated variants showed *trans*-eQTL effects on the same downstream gene. Here we observed 47 different traits where at least four independent variants affected the same gene in *trans*, 3.4× higher than expected by chance (P = 0.001; two-tailed two-sample test of equal proportions; **Supplementary Table 8**). For SLE, for example, we observed that the gene expression levels of *IFI44L, HERC5, IFI6, IFI44, RSAD2, MX1, ISG15, ANKRD55, OAS3, OAS2, OASL* and *EPSTI1* (nearly all interferon genes) were affected by at least three SLE-associated genetic variants, clearly showing the involvement of interferon signaling in SLE (Figure 5).

**Figure 5.**
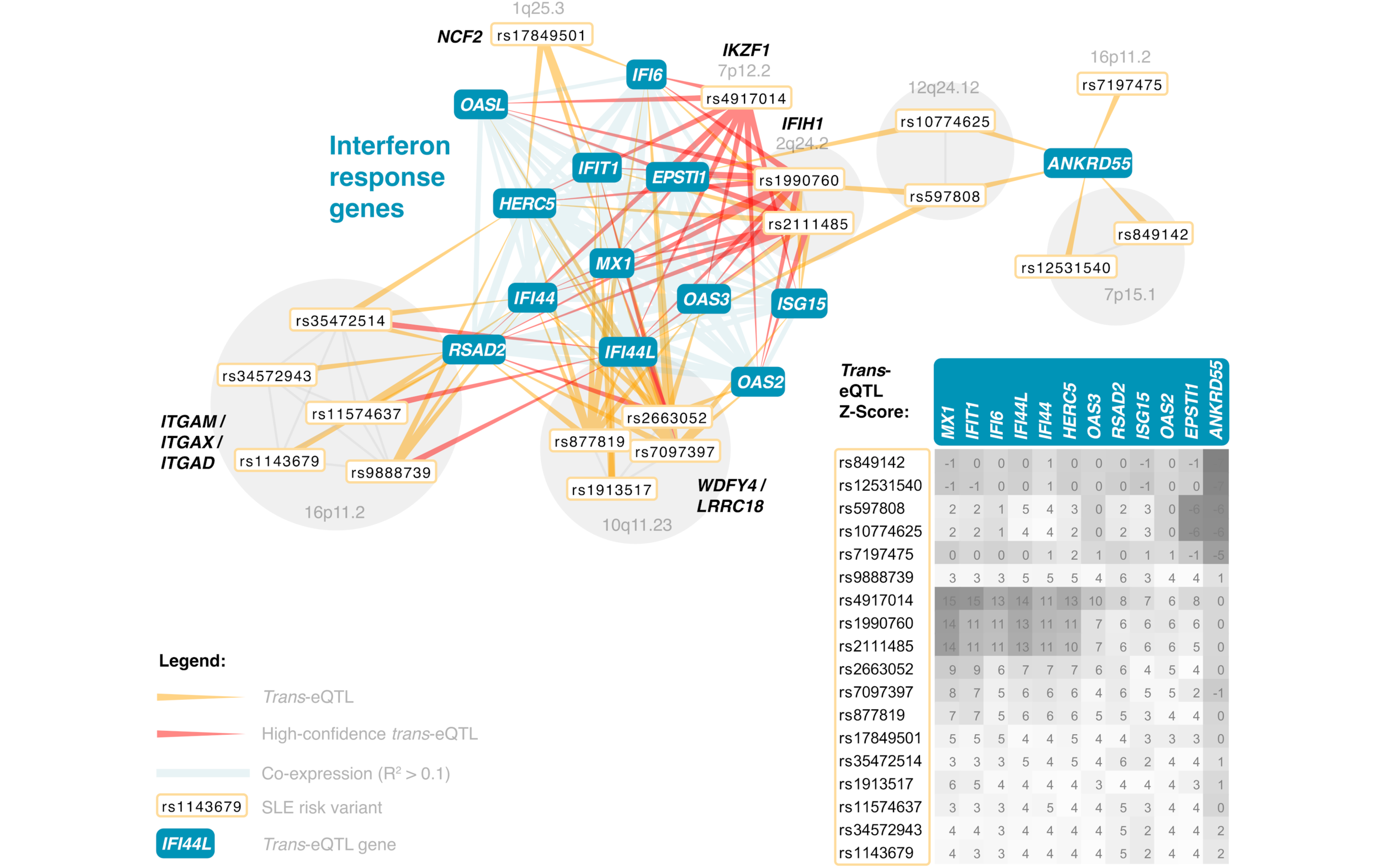
SNPs associated with SLE converge to the shared cluster of interferon response genes. Shown are genes which are affected by at least three independent GWAS SNPs. SNPs in partial LD are grouped together. Heat map indicates the direction and strength of individual *trans*-eQTL effects (Z-scores).

This convergence of multiple SNPs on the same genes lends credence to recent hypotheses with regards to the ‘omnigenic’ architecture of complex traits^8^: indeed multiple unlinked variants do affect the same ‘core’ genes. The recent omnigenic model^42^ proposes a strategy to partition between core genes, which have direct effects on a disease, and peripheral genes, which can only affect disease risk indirectly through regulation of core genes. In **Supplementary Equations**, we show that this model also implies a correlation between polygenic risk scores and expression of core genes. We therefore studied this systematically by aggregating multiple associated variants into polygenic scores and ascertaining how they correlate with gene expression levels.

## eQTSs identify key driver genes for polygenic traits

To ascertain the coordinated effects of trait-associated variants on gene expression, we used available GWAS summary statistics to calculate PGSs for 1,263 traits in 28,158 samples (**Online Methods, Supplementary Table 14**). We reasoned that when a gene shows expression levels that significantly correlate with the PGS for a specific trait (an expression quantitative trait score; eQTS), the downstream *trans*-eQTL effects of the individual risk variants converge on that gene, and hence, that the gene may be a driver of the disease.

Our meta-analysis identified 18,210 eQTS effects (FDR < 0.05), representing 689 unique traits (54%) and 2,568 unique genes (13%; **Supplementary Table 15, Figure 1A**). As expected, most eQTS associations represent blood cell traits (**Extended Data Figure 14, Supplementary Table 16):** for instance the PGS for mean corpuscular volume correlated positively with the expression levels of genes specifically expressed in erythrocytes, such as genes coding for hemoglobin subunits. However, we also identified eQTS associations for genes that are known drivers of other traits.

For example, 11 out of 26 genes associating with the PGS for high density lipoprotein levels (HDL^43,44^; FDR<0.05; **Figure 6A**) have previously been associated with lipid or cholesterol metabolism (**Supplementary Table 18**). *ABCA1* and *ABCG1*, which positively correlated with the PGS for high HDL, mediate the efflux of cholesterol from macrophage foam cells and participate in HDL formation. In macrophages, the downregulation of both *ABCA1* and *ABCG1* reduced reverse cholesterol transport into the liver by HDL^45^ (**Figure 6B**). The genetic risk for high HDL was also negatively correlated with the expression of the low density lipoprotein receptor *LDLR* (strongest eQTS P=3.35<10^-20^) known to cause hypercholesterolemia^46^. Similarly, the gene encoding the TF SREBP-2, which is known to increase the expression of *LDLR*, was downregulated (strongest eQTS P=3.08<10^-7^). The negative correlation between *SREBF2* expression and measured HDL levels has been described before^47^, indicating that the eQTS reflects an association with the actual phenotype. Zhernakova et al. proposed a model where down-regulation of *SREBF2* results in the effect on its target gene *FADS2.* We did not observe a significant HDL eQTS effect on *FADS2* (all eQTS P>0.07), possibly because the indirect effect is too small to detect. We hypothesize that HDL levels in blood can result in a stronger reverse cholesterol transport into the liver, which may result in downregulation of *LDLR*^48^

**Figure 6.**
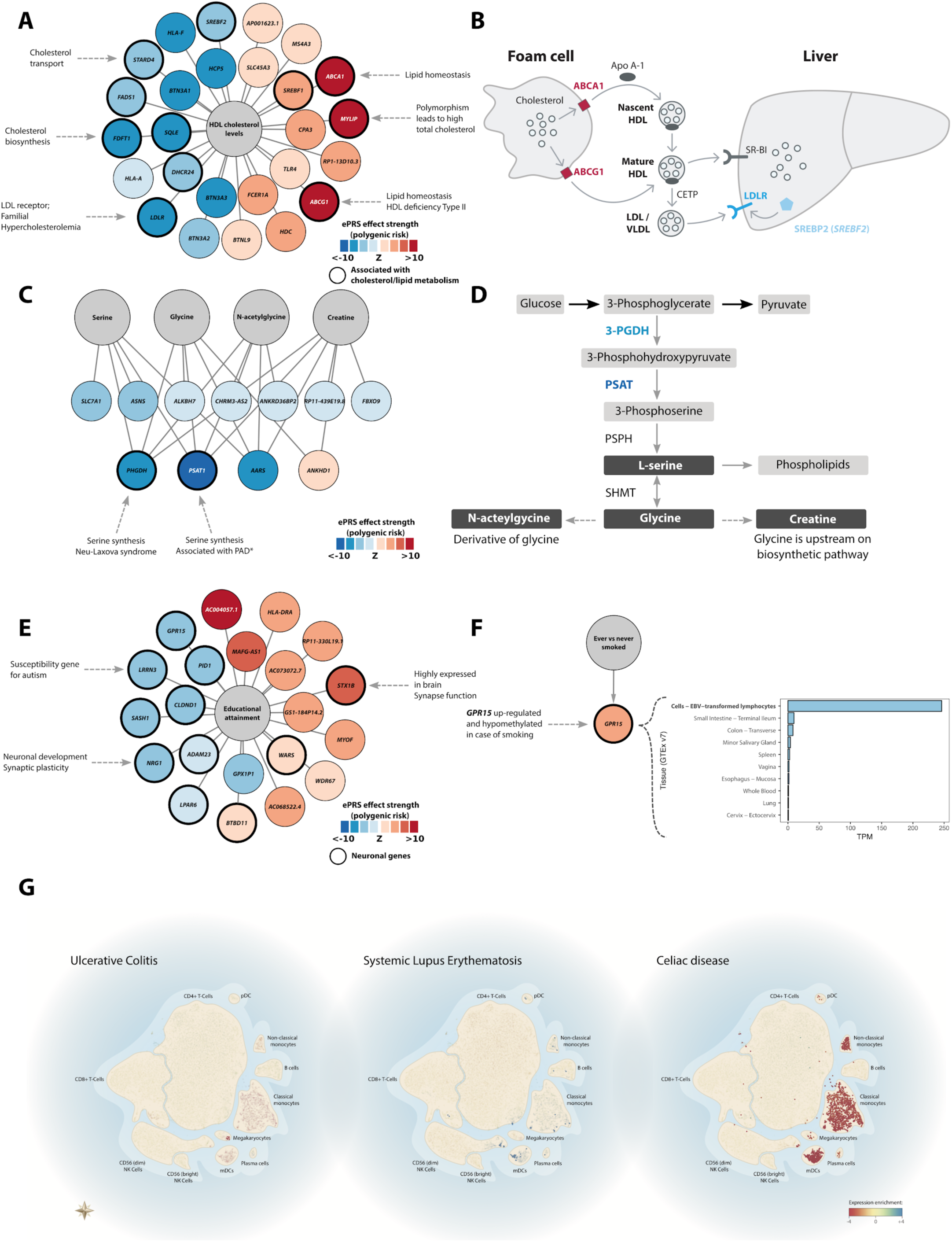
Examples of eQTS. (**A**) Polygenic risk score (PRS) for high density lipoprotein associates to lipid metabolism genes. (**B**) The role of *ABCA1, ABCG1*, and *SREBF2* in cholesterol transport. (**C**) Polygenic scores for serine, glycine, n-acetylglycine and creatine levels negatively associate with gene expression of *PHGDH, PSAT1*, and *AARS.* (**D**) Serine biosynthesis pathway. (**E**) PRS for educational attainment identifies genes with neuronal functions. (**F**) Polygenic score for smoking status upregulates *GPR15*, which plays a role in lymphocyte differentiation. (**G**) eQTS genes for immune-related diseases are enriched for genes specifically expressed in certain blood cell types.

eQTS analysis also identified genes relevant for non-blood traits, such as the association of *GPR15* (P=3.7<10^-8^, FDR<0.05; **Figure 6F**) with the trait ‘ever versus never smoking’^49^. *GPR15* is a biomarker for smoking^50^ that is overexpressed and hypomethylated in smokers^51^. We observe strong *GPR15* expression in lymphocytes (**Figure 6F**), suggesting that the association with smoking could originate from a change in the proportion of T cells in blood^52^. As *GPR15* is involved in T cell homing and has been linked to colitis and inflammatory phenotypes, it is hypothesized to play a key role in smoking-related health risks^53^.

The PGS for another non-blood trait, educational attainment^54^, correlated significantly with the expression of 21 genes (**FDR<0.05; Figure 6E, Supplementary Table 15**). Several of the strongly associated genes are known to be involved in neuronal processes (**Supplementary Table 19**) and show expression in neuronal tissues (GTEx v7, **Extended Data Figure 15**). *STX1B* (strongest eQTS P=1.3×10^-20^) is specifically expressed in brain, and its encoded protein, syntaxin 1B, participates in the exocytosis of synaptic vesicles and synaptic transmission^55^. Another gene highly expressed in brain, *LRRN3* (Leucine-rich repeat neuronal protein 3; strongest eQTS P=1.7×10^-11^) was negatively associated with the PGS for educational attainment, and has been associated with autism susceptibility^56^. The downregulated *NRG1* (neuregulin 1; strongest eQTS P=4.5×10^-7^),encodes a well-established growth factor involved in neuronal development and has been associated to synaptic plasticity^57^. *NRG1* was also positively associated with the PGS for monocyte levels^20^ (strongest eQTS P=1.5×10^-7^), several LDL cholesterol traits (e.g. medium LDL particles^44^; strongest eQTS P=6.2×10^-8^), coronary artery disease^33^ (strongest eQTS P=1.5×10^-6^) and body mass index in females^58^ (strongest eQTS P=9.2×10^-12^).

eQTS can also identify pathways known to be associated with monogenic diseases. For example, the PGSs for serine, glycine, the glycine derivative n-acetylglycine and creatine^59,60^ (**Figure 6C**) were all negatively associated with the gene expression levels of *PHGDH, PSAT1* and *AARS* (P < 5.3×10^-7^). *PHGDH* and *PSAT1* encode crucial enzymes that regulate the synthesis of serine and, in turn, glycine^61^ (**Figure 6D**), while n-acetylglycine and creatine form downstream of glycine^62^. Mutations in *PSAT1* and *PHGDH* can result in serine biosynthesis defects including phosphoserine aminotransferase deficiency^63^, phosphoglycerate dehydrogenase deficiency^64^, and Neu-Laxova syndrome^65^, all diseases characterized by low concentrations of serine and glycine in blood and severe neuronal manifestations. *AARS* encodes alanyl-tRNA synthetase, which links alanine to tRNA molecules. A mutation in *AARS* has been linked to Charcot Marie Tooth disease^66^, while the phenotypically similar hereditary sensory neuropathy type 1 (HSN1^67^) can be caused by a mutation in the gene encoding serine palmitoyltransferase. The gene facilitates serine’s role in sphingolipid metabolism^68^. Disturbances in this pathway are hypothesized to be central in the development the neuronal symptoms^69^, suggesting a link between *AARS* expression and the serine pathway. Unexpectedly, the genetic risk for higher levels of these amino acids was associated with lower expression of *PHGDH, PSAT1*, and *AARS*, implying the presence of a negative feedback loop that controls serine synthesis.

We next evaluated 6 immune diseases for which sharing of loci has been reported previously, and also observed sharing of downstream eQTS effects for these diseases (**Supplementary Table 20**). For example, the interferon gene *STAT1* was significantly associated with T1D, celiac disease (CeD), IBD and primary biliary cirrhosis (PBC). However, some of these genes are also marker genes for specific blood-cell types, such as *CD79A*, which showed a significant correlation with T1D and PBC. To test whether disease-specific eQTS gene signatures are reflected by blood cell proportions, we investigated single-cell RNA-seq data^31^ (**Online Methods; Figure 6G**). For ulcerative colitis (a subtype of IBD), we observed significant depletion of expression in megakaryocytes. SLE eQTS genes were enriched for antigen presentation (GeneNetwork P=1.3<10^-5^) and interferon signaling (GeneNetwork P=1.4<10^-4^), consistent with the well-described interferon signature in SLE patients^70,71^. Moreover, the SLE genes were significantly enriched for expression in mature dendritic cells, whose maturation depends on interferon signaling^72^. For CeD, we observed strong depletion of eQTS genes in monocytes and dendritic cells, and a slight enrichment in CD4+ and CD8+ T cells. The enrichment of cytokine (GeneNetwork P=1.6<10^-15^) and interferon (GeneNetwork P=7.8<10^-13^) signaling among the CeD eQTS genes is expected as a result of increased T cell populations.

## Cell-type-specificity of eQTS associations

We next ascertained to what extent these eQTS associations can be replicated in non-blood tissues. We therefore aimed to replicate the significant eQTS effects in 1,460 LCL samples and 762 iPSC samples. Due to the fact these cohorts have a comparatively low sample sizes and study different cell types, we observed limited replication: 10 eQTS showed significant replication effect (FDR<0.05) in the LCL dataset, with 9 out of those (90%) showing the same effect direction as in the discovery set (**Extended Data Figure 16A, Supplementary Table 17**). For iPSCs, only 5 eQTS showed a significant effect (**Extended Data Figure 16B, Supplementary Table 17**). Since only a few eQTS associations are significant in non-blood tissues and the majority of identified eQTS associations are for blood-related traits, we speculate these effects are likely to be highly cell-type specific. This indicates that large-scale eQTL meta-analyses in other tissues could uncover more genes on which trait-associated SNPs converge.

## Discussion

We here performed *cis*-eQTL, *trans*-eQTL and eQTS analyses in 31,684 blood samples, reflecting a six-fold increase over earlier large-scale studies^1,5^. We identified *cis*-eQTL effects for 88.3% and *trans*-eQTL effects for 32% of all genes that are expressed in blood.

We observed that *cis*-eQTL SNPs map close to the TSS or TES of the *cis*-gene: for the top 20% strongest *cis*-eQTL genes, 84.1% of the lead eQTL SNPs map within 20kb of the gene, indicating that these are variants immediately adjacent to the start or end of transcripts that primarily drive *cis*-eQTL effects. The trait-associated variants that we studied showed a different pattern: 77.4% map within 20kb of the closest protein-coding gene, suggesting that the genetic architecture of *cis*-eQTLs is different from disease-associated variants. This is supported by the epigenetic differences that we observed between these two groups and can also partly explain, why we did not observe significantly increased overlap between genes prioritized using pathway enrichment analysis^15^ and genes prioritized using our *cis*-eQTLs. In contrast, for numerous traits we observed that multiple unlinked *trans*-eQTL variants often converge on genes with a known role in the biology for these traits (e.g. the involvement of interferon genes in SLE).

We therefore focused on *trans*-eQTL and eQTS results to gain insight into trait-relevant genes and pathways (**Figures 4, 6**). We estimate that 17.4% of our *trans*-eQTLs are driven by transcriptional regulation, whereas the remaining fraction is driven by not-yet-identified mechanisms. Our results support a model which postulates that, compared to *cis*-eQTLs, weaker distal and polygenic effects converge on core (key driver) genes that are more relevant to the traits and more specific for trait-relevant cell types (**Figure 1B**). The examples we have highlighted demonstrate how insights can be gained from our resource, and we envision similar interpretation strategies can be applied to the other identified *trans*-eQTL and eQTS effects. The catalog of genetic effects on gene expression we present here (available at http://www.eqtlgen.org) is a unique compendium for the development and application of novel methods that prioritize causal genes for complex traits^14 73^, as well as for interpreting the results of genome-wide association studies.

## Methods

### Cohorts

EQTLGen Consortium data consists of 31,684 blood and PBMC samples from 37 datasets, preprocessed in a standardized way and analyzed by each cohort analyst using the same settings (**Online Methods**). 26,886 (85%) of the samples added to discovery analysis were whole blood samples and 4,798 (15%) were PBMCs, and the majority of samples were of European ancestry (**Supplementary Table 1**). The gene expression levels of the samples were profiled by Illumina (N=17,421; 55%), Affymetrix U291 (N=2,767; 8.7%), Affymetrix HuEx v1.0 ST (N=5,075; 16%) expression arrays and by RNA-seq (N=6,422; 20.3%). A summary of each dataset is outlined in **Supplementary Table 1**. Detailed cohort descriptions can be found in the **Supplementary Note**. Each of the cohorts completed genotype and expression data pre-processing, PGS calculation, *cis*-eQTL-, *trans*-eQTL and eQTS-mapping, following the steps outlined in the online analysis plans, specific for each platform (see **URL**s) or with slight alterations as described in **Supplementary Table 1** and the **Supplementary Note**. All but one cohort (Framingham Heart Study), included non-related individuals into the analysis.

### Genotype data preprocessing

The primary pre-processing and quality control of genotype data was conducted by each cohort, as specified in the original publications and in the **Supplementary Note**. The majority of cohorts used genotypes imputed to 1kG p1v3 or a newer reference panel. GenotypeHarmonizer^74^ was used to harmonize all genotype datasets to match the GIANT 1kG p1v3 ALL reference panel and to fix potential strand issues for A/T and C/G SNPs. Each cohort tested SNPs with minor allele frequency (MAF) > 0.01, Hardy-Weinberg P-value > 0.0001, call rate > 0.95, and MACH r^2^ > 0.5.

## Expression data preprocessing

### Illumina array

Illumina array datasets expression were profiled by HT-12v3, HT-12v4 and HT-12v4 WGDASL arrays. Before analysis, all the probe sequences from the manifest files of those platforms were remapped to GRCh37.p10 human genome and transcriptome, using SHRiMP v2.2.3 aligner^75^ and allowing 2 mismatches. Probes mapping to multiple locations in the genome were removed from further analyses.

For Illumina arrays, the raw unprocessed expression matrix was exported from GenomeStudio. Before any pre-processing, the first two principal components (PCs) were calculated on the expression data and plotted to identify and exclude outlier samples. The data was normalized in several steps: quantile normalization, log_2_ normalization, probe centering and scaling by the equation Expression_Probe,Sample_ = (Expression_Probe,Sample_ - MeanProbe) / Std.Dev._probe_. Genes showing no variance were removed. Next, the first four multidimensional scaling (MDS) components, calculated based on non-imputed and pruned genotypes using plink v1.07^76^, were regressed out of the expression matrix to account for population stratification. We further removed up to 20 first expression-based PCs that were not associated to any SNPs, as these capture non-genetic variation in expression. Each cohort also ran MixupMapper^77^ software to identify incorrectly labeled genotype-expression combinations, and to remove identified sample mix-ups.

### Affymetrix arrays

Affymetrix array-based datasets used the expression data previously pre-processed and quality controlled as indicated in the **Supplementary Note**.

### RNA-seq

Alignment, initial quality control and quantification differed slightly across datasets, as described in the **Supplementary Note**. Each cohort removed outliers as described above, and then used Trimmed Mean of M-values (TMM) normalization and a counts per million (CPM) filter to include genes with >0.5 CPM in at least 1% of the samples. Other steps were identical to Illumina processing, with some exceptions for the BIOS Consortium datasets (**Supplementary Note**).

## *cis*-eQTL mapping

*cis*-eQTL mapping was performed in each cohort using a pipeline described previously^1^. In brief, the pipeline takes a window of 1Mb upstream and 1Mb downstream around each SNP to select genes or expression probes to test, based on the center position of the gene or probe. The associations between these SNP-gene combinations was calculated using a Spearman correlation. Next, 10 permutation rounds were performed by shuffling the links between genotype and expression identifiers and re-calculating associations. The false discovery rate (FDR) was determined using 10 meta-analyzed permutations: for each gene in the real analysis, the most significant association was recorded, and the same was done for each of the permutations, resulting in a gene-level FDR. *cis*-eQTLs with a gene-level FDR < 0.05 (corresponding to P < 1.829<10^-5^) and tested in at least two cohorts were deemed significant.

## *Trans-eQTL* mapping

*Trans*-eQTL mapping was performed using a previously described pipeline^1^ while testing a subset of 10,317 SNPs previously associated with complex traits. We required the distance between the SNP and the center of the gene or probe to be >5Mb. To maximize the power to identify *trans*-eQTL effects, the results of the summary statistics based or iterative conditional *cis*-eQTL mapping analyses (**Supplementary Note**) were used to correct the expression matrices before *trans*-eQTL mapping. For that, top SNPs for significant conditional *cis*-eQTLs were regressed out from the expression matrix. Finally, we removed potential false positive *trans*-eQTLs caused by reads crossmapping with *cis* regions (**Supplementary Note**).

### Genetic risk factor selection

Genetic risk factors were downloaded from three public repositories: the EBI GWAS Catalogue^78^ (downloaded 21.11.2016), the NIH GWAS Catalogue and Immunobase (www.immunobase.org;accessed26.04.2016), applying a significance threshold of P ≤ 5<10^-8^. Additionally, we added 2,706 genome-wide significant GWAS SNPs from a recent blood trait GWAS^20^. SNP coordinates were lifted to hg19 using the *liftOver* command from R package rtracklayer v1.34.1^79^ and subsequently standardized to match the GIANT 1kG p1v3 ALL reference panel. This yielded 10,562 SNPs (**Supplementary Table 2**). We tested associations between all risk factors and genes that were at least 5Mb away to ensure that that they did not tag a *cis*-eQTL effect. All together, 10,317 trait-associated SNPs were tested in *trans*-eQTL analyses.

## eQTS mapping

### PGS trait inclusion

Full association summary statistics were downloaded from several publicly available resources (**Supplementary Table 13**). GWAS performed exclusively in non-European cohorts were omitted. Filters applied to the separate data sources are indicated in the **Supplementary Note**. All the dbSNP rs numbers were standardized to match GIANT 1kG p1v3, and the directions of effects were standardized to correspond to the GIANT 1kG p1v3 minor allele. SNPs with opposite alleles compared to GIANT alleles were flipped. SNPs with A/T and C/G alleles, tri-allelic SNPs, indels, SNPs with different alleles in GIANT 1kG p1v3 and SNPs with unknown alleles were removed from the analysis. Genomic control was applied to all the P-values for the datasets not genotyped by Immunochip or Metabochip. Additionally, genomic control was skipped for one dataset that did not have full associations available^80^ and for all the datasets from the GIANT consortium, as for these genomic control had already been applied. All together, 1,263 summary statistic files were added to the analysis. Information about the summary statistics files can be found in the **Supplementary Note** and **Supplementary Table 14**.

### PGS calculation

A custom Java program, GeneticRiskScoreCalculator-v0.1.0c, was used for calculating several PGS in parallel. Independent effect SNPs for each summary statistics file were identified by double-clumping by first using a 250kb window and subsequently a 10Mb window with LD threshold R^2^=0.1. Subsequently, weighted PGS were calculated by summing the risk alleles for each independent SNP, weighted by its GWAS effect size (beta or log(OR) from the GWAS study). Four GWAS P-value thresholds (P<5×10^-8^, 1<10^-5^, 1×10^-4^ and 1×10^-3^) were used for constructing PGS for each summary statistics file.

### Pruning the SNPs and PGS

To identify a set of independent genetic risk factors, we conducted LD-based pruning as implemented in PLINK 1.9^81^ with the setting-indep-pairwise 50 5 0.1. This yielded in 4,586 uncorrelated SNPs (R^2^<0.1, GIANT 1kG p1v3 ALL).

To identify the set of uncorrelated PGS, ten permuted *trans*-eQTL Z-score matrices from the combined *trans*-eQTL analysis were first confined to the pruned set of SNPs. Those matrices were then used to identify 3,042 uncorrelated genes, based on Z-score correlations (absolute Pearson R <0.05). Next, permuted eQTS Z-score matrices were confined to uncorrelated genes and used to calculate pairwise correlations between all genetic risk scores to define a set of 1,873 uncorrelated genetic risk scores (Pearson R^2^ < 0.1).

## Empirical probe matching

To integrate different expression platforms (four different Illumina array models, RNA-seq, Affymetrix U291 and Affymetrix Hu-Ex v1.0 ST) for the purpose of meta-analysis, we developed an empirical probe-matching approach. We used the pruned set of SNPs to conduct per-platform meta-analyses for all Illumina arrays, for all RNA-seq datasets, and for each Affymetrix dataset separately, using summary statistics from analyses without any gene expression correction for principal components. For each platform, this yielded an empirical *trans*-eQTL Z-score matrix, as well as ten permuted Z-score matrices, where links between genotype and expression files were permuted. Those permuted Z-score matrices reflect the gene-gene or probe-probe correlation structure.

We used RNA-seq permuted Z-score matrices as a gold standard reference and calculated for each gene the Pearson correlation coefficients with all the other genes, yielding a correlation profile for each gene. We then repeated the same analysis for the Illumina meta-analysis, and the two different Affymetrix platforms. Finally, we correlated the correlation profiles from each array platform with the correlation profiles from RNA-seq. For each array platform, we selected the probe showing the highest Pearson correlation with the corresponding gene in the RNA-seq data and treated those as matching expression features in the combined meta-analyses. This yielded 19,960 genes that were detected in RNA-seq datasets and tested in the combined meta-analyses. Genes and probes were matched to Ensembl v71^82^ (see **URL**s) stable gene IDs and HGNC symbols in all the analyses.

## Cross-platform replications

To test the performance of the empirical probe-matching approach, we conducted discovery *cis*-, *trans-* and eQTS meta-analyses for each expression platform (RNA-seq, Illumina, Affymetrix U291 and Affymetrix Hu-Ex v1.0 ST arrays; array probes matched to 19,960 genes by empirical probe matching). For each discovery analysis, we conducted replication analyses in the three remaining platforms, observing strong replication of both *cis*-eQTLs, *trans*-eQTLs and eQTS in different platforms, with very good concordance in allelic direction.

## Meta-analyses

We meta-analyzed the results using a weighted Z-score method^1^, where the Z-scores are weighted by the square root of the sample size of the cohort. For *cis*-eQTL and *trans*-eQTL meta-analyses, this resulted in a final sample size of N=31,684. The combined eQTS meta-analysis included the subset of unrelated individuals from the Framingham Heart Study, resulting in a combined sample size of 28,158.

## Quality control of the meta-analyses

For quality control of the overall meta-analysis results, MAFs for all tested SNPs were compared between eQTLGen and 1kG p1v3 EUR (**Extended Data Figure 3**), and the effect direction of each dataset was compared against the meta-analyzed effect (**Extended Data Figure 2A-C**).

## FDR calculation for *trans*-eQTL and eQTS mapping

To determine nominal P-value thresholds corresponding to FDR=0.05, we used the pruned set of SNPs for *trans*-eQTL mapping and permutation-based FDR calculation, as described previously^1^. We leveraged those results to determine the P-value threshold corresponding to FDR=0.05 and used this as a significance level in *trans*-eQTL mapping in which all 10,317 genetic trait-associated SNPs were tested. In the eQTS analysis, an analogous FDR calculation was performed using a pruned set of PGSs. We analyzed only SNP/PGS-gene pairs tested in at least two cohorts.

## Positive and negative set of *trans*-eQTLs

Based on the results of integrative *trans*-eQTL mapping, we defined true positive (TP) and true negative (TN) sets of *trans*-eQTLs. TP set was considered as all significant (FDR<0.05) *trans*-eQTLs. TN set of *trans*-eQTLs was selected as non-significant (max absolute meta-analysis Z-score 3; all FDR>0.05) SNP-gene combinations, adhering to following conditions:

1. The size of TN set was set equal to the size of TP set (59,786 *trans*-eQTLs).
2. Each SNP giving *trans*-eQTL effects on X genes in the TP set, is also giving *trans*-eQTL effects on X genes in the TN set.
3. Each gene that is affected in *trans* by Y SNPs in the TP set, is also affected in *trans* by Y SNPs in the TN set.
4. Adhere to the correlation structure of the SNPs: if two SNPs are in perfect LD, they affect the same set of genes, both in the TP set and in the TN set.
5. Adhere to the correlation structure of the genes: if two genes are perfectly co-expressed, they are affected by the same SNPs, both in the TP set and in the TN set.

This set of TN *trans*-eQTLs was used in subsequent enrichment analyses as the matching set for comparison.

## Conditional *trans*-eQTL analyses

We aimed to estimate how many *trans*-eQTL SNPs were likely to drive both the *trans*-eQTL effect and the GWAS phenotype. The workflow of this analysis is shown in **Extended Data Figure 6**. We used the integrative *trans*-eQTL analysis results as an input, confined ourselves to those effects which were present in the datasets we had direct access to (BBMRI-BIOS+EGCUT; N=4,339), and showed nominal P<8.3115× 10^-06^ in the meta-analysis of those datasets. This P-value threshold was the same as in the full combined *trans*-eQTL meta-analysis and was based on the FDR=0.05 significance threshold identified from the analysis run on the pruned set of GWAS SNPs after removal of cross-mapping effects. We used the same methods and SNP filters as in the full combined *trans*-eQTL meta-analysis, aside from the FDR calculation, which was based on the full set of SNPs, instead of the pruned set of SNPs.

For each significant *trans*-eQTL SNP, we defined the locus by adding a ±1Mb window around it. Next, for each *trans*-eQTL gene we ran iterative conditional *trans*-eQTL analysis using all loci for given *trans*-eQTL gene. We then evaluated the LD between all conditional top *trans*-eQTL SNPs and GWAS SNPs using a 1 Mb window and R^2^>0.8 (1kG p1v3 EUR) as a threshold for LD overlap.

## *Trans*-eQTL mediation analysis

To identify potential mediators of *trans*-eQTL effects we used a G x E interaction model:

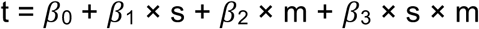

Where t is the expression of the *trans*-eQTL gene, s is the *trans*-eQTL SNP, and m is the expression of a potential mediator gene within 100kb of the *trans*-eQTL SNP. On top of the gene expression normalization that we used for the rest of our analysis, we used a rank-based inverse normal transformation to enforce a normal distribution before fitting the linear model, identical to the normalization used by Zhernakova et al.^47^ in their G × E interaction eQTL analyses. We fitted this model separately on each of the cohorts that are part of the BIOS consortium. We transformed the interaction P-values to Z-scores and used the weighted Z-score method^83^ to perform a meta-analysis on the in total 3,831 samples. The Benjamini & Hochberg procedure^84^ was used to limit the FDR to 0.05. The plots in **Extended Data Figure 9** are created with the default normalization, the regression lines are the best-fitting lines between the mediator gene and the *trans* eQTL gene,stratified by genotype. We used a Fisher's exact test to calculate the enrichment of significant (FDR≤0.05) interactions between our TP *trans*-eQTLs and the interactions identified in the TN *trans-* eQTL set.

## TF and tissue enrichment analyses

We downloaded the curated sets of known TF targets and tissue-expressed genes from the Enrichr web site^85,86^. TF target gene sets included TF targets as assayed by ChIP-X experiments from ChEA^87^ and ENCODE^88,89^ projects, and tissue-expressed genes were based on the ARCHS4 database^90^. Those gene sets were used to conduct hypergeometric over-representation analyses as implemented into the R package ClusterProfiler^91^.

## SMR analyses

To gain further insight into genes that are important in the biology of the trait, we used the combined *cis*-eQTL results to perform SMR^14^ for 16 large GWAS studies (**Supplementary Table 20**). We derived *cis*-eQTL beta and standard error of the beta (SE(beta)) from the Z-score and the MAF reported in 1kG v1p3 ALL, using the following formulae^14^

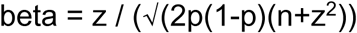

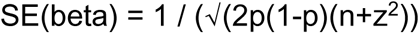

Where p is the MAF and n is the sample size.

The *cis*-eQTLs were converted to the dense BESD format. The 1 kG p1v3 ALL reference panel was also used to calculate LD, and SMR analysis was run using the SMR software v0.706 without any P-value cut-offs on either GWAS or eQTL input.

## DEPICT

We applied DEPICT v194^15^ to the same 16 recent GWAS traits as above (**Supplementary Table 20**), using all variants that attain a genome-wide significant P-value threshold. Specifically, we looked at the gene prioritization and gene set enrichment analyses to compare the results with the output of other prioritization methods (SMR ^14^).

## Comparison of gene prioritization with DEPICT and SMR

To investigate the consistency between results from two gene prioritization methods, we compared the enrichment of overlapping genes for 16 GWAS traits (**Supplementary Table 20**). We confined ourselves to genes that were tested in SMR and that fell within the DEPICT loci, and tested whether genes significant in SMR (P-value < 0.05 / number of tested genes) and DEPICT (FDR < 0.05) were enriched (one-sided Fisher’s Exact Test).

## Epigenetic marks enrichment

We ascertained epigenetic properties of the lead *cis*-eQTL SNPs, and contrasted these to a set of 3,688 trait-associated SNPs that were associated with either blood-related traits (such as mean corpuscular volume or platelet counts) or immune-mediated diseases. The SNPs were annotated with histone and chromatin marks information from the Epigenomics Roadmap Project. We summarized the information by calculating the overlap ratio across 127 human cell types between the epigenetic marks and the SNP within a window size of +/-25bp: if a SNP co-localizes with a mark for all 127 cell-types, the score for that SNP will be 1; if a SNP co-localizes with a mark for none of the cell-types, the score will be 0.

The reason we chose only SNPs associated to blood-related traits and immune-mediated diseases was to minimize potential confounding due to a subtle bias in the Epigenomics Roadmap Project towards blood cell-types: 29 of the 127 cell-types that we studied were blood cell types. However, when redoing the epigenetic enrichment analysis, while excluding these blood cell types, we did not see substantial differences in the enriched and depleted histone marks.

## Chromosomal contact analyses

### Capture Hi-C overlap for *cis*-eQTLs

To assess whether *cis*-eQTL lead SNPs overlapped with chromosomal contact as measured using Hi-C data, we used promoter capture Hi-C data^92^, downloaded from CHiCP^93^ (see URLs). We took the lead eQTL SNPs and overlapped these with the capture Hi-C data and studied the 10,428 cis-eQTL genes for which this data is available. We then checked whether the Capture Hi-C target maps within 5kb of the lead SNP. Of 508 *cis*-eQTL genes that mapped over 100 kb from the TSS or TES, 223 overlapped capture Hi-C data (27.8%). Of 7,984 *cis*-eQTL genes that mapped within 100kb from the TSS or TES, 1,641 overlapped capture Hi-C data (17.0%, Chi^2^ test P = 10^-14^). To ensure this was not an artefact, we performed the same analysis, while flipping the location of the capture Hi-C target with respect to the location of the bait, and did not observe any significant difference (Chi^2^ test P = 0.59).

### Hi-C overlap enrichment analysis for *trans-eQTLs*

To assess whether *trans*-eQTLs were enriched for chromosomal contacts as measured using HiC data, we downloaded the contact matrices for the human lymphoblastoid GM12878 cell line^19^ (GEO accession GSE63525). We used the intrachromosomal data at a resolution of 10kb with mapping quality of 30 or more (MAPQGE30), and normalized using the KRnorm vectors. For each of the 59,786 *trans*-eQTLs, we evaluated whether any contact was reported in this dataset. We divided each *trans*-eQTL SNP and any of their proxies (R^2^>0.8, 1kG p1v3, EUR, acquired from SNiPA^94^; **URL**s) in 10kb blocks. The *trans*-eQTL genes were also assigned to 10kb blocks, and to multiple blocks if the gene was more than 10kb in length (length between TSS and TES, Ensembl v71). For each individual *trans*-eQTL SNP-gene pair, we then determined if there was any overlap with the Hi-C contact matrices. We repeated this analysis using the true negative set of *trans-eQTLs* described before to generate a background distribution of expected contact.

## Data availability

Full summary statistics from eQTLGen meta-analyses are available on the eQTLGen website: www.eqtlgen.org which was built using the MOLGENIS framework^95^.

## Code availability

Individual cohorts participating in the study followed the analysis plans as specified in the **URL**s or with slight alterations as described in the **Methods** and **Supplementary Note**. All tools and source code, used for genotype harmonization, identification of sample mixups, eQTL mapping, meta-analyses and for calculating polygenic scores are freely available at https://github.com/molgenis/systemsgenetics/.

## Acknowledgments

The cohorts participating in this study list the acknowledgments in the cohort-specific supplemental information in Supplementary Note.

We thank i2QTL CONSORTIUM for providing the iPSC replication results.

We thank Kate McIntyre for editing the final text.

This work is supported by a grant from the European Research Council (ERC Starting Grant agreement number 637640 ImmRisk) to LF and a VIDI grant (917.14.374) from the Netherlands Organisation for Scientific Research (NWO) to LF. We thank the UMCG Genomics Coordination Center, MOLGENIS team, the UG Center for Information Technology, and the UMCG research IT program and their sponsors in particular BBMRI-NL for data storage, high performance compute and web hosting infrastructure. BBMRI-NL is a research infrastructure financed by the Netherlands Organization for Scientific Research (NWO) [grant number 184.033.111].

## URLs

Full summary statistics from this study, www.eqtlgen.org

ExAC pLI scores, http://exac.broadinstitute.org/downloads;

Ensembl v71 annotation file, ftp://ftp.ensembl.org/pub/release-71/gtf/homosapiens;

Reference for genotype harmonizing, ftp://share.sph.umich.edu/1000genomes/fullProject/2012.03.14/GIANT.phase1_release_v3.20101123.snps_indels_svs.genotypes.refpanel.ALL.vcf.gz.tgz

EQTLGen analysis plan for Illumina array datasets, https://github.com/molgenis/systemsgenetics/wiki/eQTL-mapping-analysis-cookbook;

EQTLGen analysis plan for RNA-seq datasets, https://github.com/molgenis/systemsgenetics/wiki/eQTL-mapping-analysis-cookbook-for-RNA-seq-data;

EQTLGen analysis plan for Affymetrix array datasets, https://github.com/molgenis/systemsgenetics/wiki/QTL-mapping-analysis-cookbook-for-Affymetrix-expression-arrays;

GenotypeHarmonizer, https://github.com/molgenis/systemsgenetics/wiki/Genotype-Harmonizer;

Protocol to resolve sample mixups, https://github.com/molgenis/systemsgenetics/wiki/Resolving-mixups;

Enrichr gene set enrichment libraries, http://amp.pharm.mssm.edu/Enrichr/;

GeneOverlap package for enrichment analyses, https://www.bioconductor.org/packages/release/bioc/html/GeneOverlap.html;

SHRiMP aligner used for re-mapping Illumina probes, http://compbio.cs.toronto.edu/shrimp/;

EBI GWAS Catalogue, https://www.ebi.ac.uk/gwas/;

Immunobase, http://www.immunobase.org/;

ClusterProfiler package used for tissue enrichment analyses, http://bioconductor.org/packages/release/bioc/html/clusterProfiler.html;

Capture Hi-C data, https://www.chicp.org/

SNiPA, used to acquire proxy SNPs, http://snipa.helmholtz-muenchen.de/snipa3/

Regulatory Circuits, used to acquire TF data, www.RegulatoryCircuits.org

